# Quantitative Pharmacology Methods for Bispecific T Cell Engagers

**DOI:** 10.1101/2025.04.23.650214

**Authors:** Mahdiar Sadeghi, Irina Kareva, Gleb Pogudin, Eduardo D. Sontag

## Abstract

T Cell Engager (TCE)s are an exciting therapeutic modality in immuno-oncology that acts to bypass antigen presentation and forms a direct link between cancer and immune cells in the Tumor Microenvironment (TME). TCEs are efficacious only when the drug is bound to both immune and cancer cell targets. Therefore, approaches that maximize the formation of the drug-target trimer in the TME are expected to increase the drug’s efficacy. In this study, we quantitatively investigate how the concentration of ternary complex and its biodistribution depend on both the targets’ specific properties and the design characteristics of the TCE, and specifically on the binding kinetics of the drug to its targets. A simplified mathematical model of drug-target interactions is considered here, with insights from the “three-body” problem applied to the model. Parameter identifiability analysis performed on the model demonstrates that steady state data, which is often available at the early pre-clinical stages, is sufficient to estimate the binding affinity of the TCE molecule to both targets. We used the model to analyze several existing antibodies, both clinically approved and under development, to explore their common kinetic features. The manuscript concludes with an assessment of a full quantitative pharmacology model that accounts for drug disposition into the peripheral compartment.

## 1 Introduction

Cancer immunotherapy has revolutionized the field of cancer treatment, highlighting that the immune system can eliminate tumors in many cases if given assistance [1]. Immunotherapy approaches include reactivating immune cells through checkpoint inhibition [2], as well as through exogenous immune cell therapies such as using Chimeric Antigen Receptor T (CAR-T) cells [3]. Despite these advances, only a subset of cancers responds to these therapies (“hot” tumors), with some tumors remaining “cold”, possibly because of reduced immune infiltration or lack of effective antigen presentation in the TME [4, 5].

One approach to circumvent this issue is to bypass antigen presentation altogether, connecting a therapeutic agent directly with a tumor cell, a strategy implemented using bispecific T cell engagers, or TCEs [6, 7, 8, 9]. This therapeutic modality has been proposed for treating acute myeloid leukemia [10], multiple myeloma [11], lymphoblastic leukemia [12], and refractory solid tumors [13]. TCE is a promising *platform* for targeted therapy across different tumor types [14]. A list of TCE molecules that were considered in this study is presented in Table 1. A more detailed review of the existing TCEs can be found in [9, 15].

**Table 1:** Existing CD3-based T Cell Engagers, sorted by the year of the referenced publication. TCEs might be engineered in different antibody structures, e.g. TCE, Half-Life Extended Bispecific T Cell Engager (HLE-TCE), and Knob-into-Holes (KIH). The targeted receptor protein of immune cells is CD3 for all the molecules considered in this study.

| TCE Molecule | Target | $K_{D1}$(nM) | $K_{D2}$(nM) | Type | Structure | kDa | Ref |
| --- | --- | --- | --- | --- | --- | --- | --- |
| Blinatumomab | CD19 | $\sim 100$ | $\sim 1$ | Liquid | TCE | 54 | [20] |
| Solitumab | Ep-CAM | $16 \pm 12$ | $77 \pm 39$ | Liquid | TCE | 55 | [21] |
| BAY2010112 | PSMA | $9.4 \pm 4.3$ | $47.0 \pm 8.1$ | Solid | TCE | 55 | [22] |
| PF-06671008 | Pcad | $11.5 \pm 0.9$ | $0.521 \pm 0.162$ | Solid | DART | 57 | [23] |
| CD3<sub>ε</sub>L/HER2 | HER2 | 50 | 0.5 | Solid | KIH | > 100 | [24] |
| CD3<sub>ε</sub>H/HER2 | HER2 | 0.5 | 0.5 | Solid | KIH | > 100 | [24] |
| CD3<sub>ε</sub>VH/HER2 | HER2 | 0.05 | 0.5 | Solid | KIH | > 100 | [24] |
| 7370 | FLT3 | 27 | $49 \times 10^{-3}$ | Liquid | IgG-based | 146 | [25] |
| PF-07062119 | GUCY2C | $7.47 \pm 0.15$ | $23.97 \pm 0.97$ | Solid | IgG1-FcyR | > 100 | [26] |
| Tarlatamab | DLL3 | $0.64 \pm 0.05$ | $14.9 \pm 0.4$ | Solid | HLE-TCE | > 100 | [27] |
| Acapatamab | PSMA | $22.4 \pm 2.8$ | $14.8 \pm 2$ | Solid | HLE-TCE | > 100 | [28] |
| Anti-CD79b/CD3 | CD79b | 12.8 | 1 | Liquid | KIH | 150 | [29] |

In this study, the first objective is to evaluate the design characteristics of existing TCEs based on the simple model presented in Figure 1. The TCE drug, *X*, targets receptors on immune cells, *T*_1_, which for TCEs is often CD3 or CD28 [16, 17], and receptors on cancer cells, *T*_2_. The drug can bind reversibly to either target, forming *X*-*T*_1_ dimer *D*_1_ at a rate constant 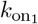 and dissociating at a rate constant 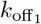, or forming the *X*-*T*_2_ dimer *D*_2_ at a rate constant 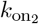 and dissociating at a rate constant 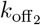. Finally, when either of the dimers bind to the remaining free target, they can form a trimer *Y*, which is the *T*_1_-*X*-*T*_2_ ternary complex. We call this system, involving immune cells, drug, and cancer cells, a “three-body” system. The key objective of this study is to find binding properties for the drug *X* on either arm given the properties of the other two targets, in such a way as to maximize concentration of *Y*. A schematic diagram of this process is given in Figure 1(a).

**Figure 1:**
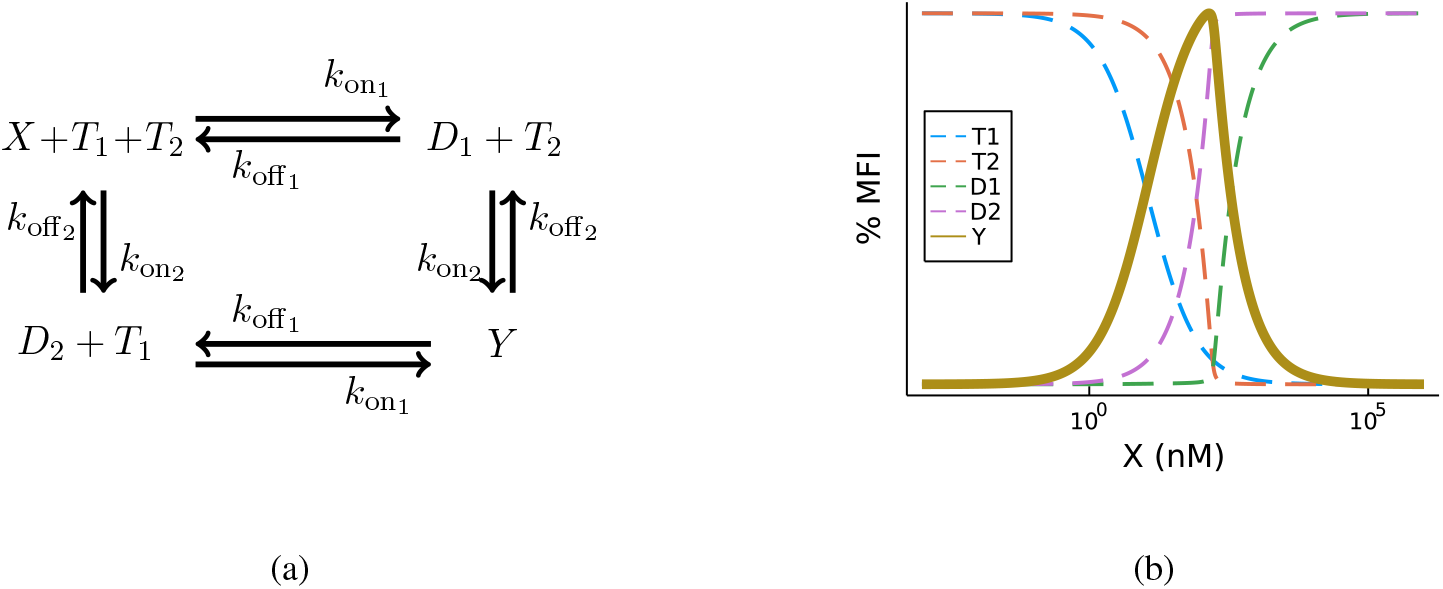
A schematic of the “three-body” model and the nominal *bell-shape* pattern. On the left side (a), the three-body model described in terms of TCE, where *X* is the initial drug concentration, *T*_1_ is the target receptor expressed on immune cells, and *T*_2_ is the target receptors expressed on cancer cells. The drug can reversibly bind to either target to form dimers *D*_1_ and *D*_2_ or bind to both to form trimer complex *Y*. Maximizing *Y* is expected to maximize drug efficacy. On the right side (b), a nominal bell-shape pattern of the ternary complex *Y* normalized to Maximum Fluorescent Intensity (MFI) at steady state is visualized as a function of initial TCE drug concentration (X in nM). Binding kinetics and initial conditions are adopted from [19]. The chemical reactions are visualized in a simple form in order to provide an intuitive idea of the reactions, and the diagram does not follow the conventions of a chemical reaction network: specifically, *T*_2_ (*T*_1_) is not involved in the dimerization reaction between *X* and *T*_1_(*T*_2_).

One particular challenge with these types of drugs is that the efficacy curve for TCEs is bell-shaped, rather than the standard Emax curve. By “bell shape”, we mean a nonmonotonic function with a single maximum at some intermediate concentration, whereas the Emax curve is a typical pharmacology model that is monotonic and saturable [18]. A TCE drug efficacy is maximized only when the drug is bound to both targets (Figure 1(b), where Y exhibits a bell-shaped curve); dimers in this drug construct (Figure 1(b), where *D*_1_ and *D*_2_ exhibit an Emax curve) are not expected to exhibit efficacy. This creates an important challenge, since, while for a typical Emax curve, higher drug concentration may result in an efficacy plateau, and increase in dose is mostly likely to just increase toxicity. For a bell-shaped efficacy curve (Figure 1(b), yellow thick line), increase in drug concentration will result not only in higher toxicity but also in loss of efficacy. As such, estimating maximally efficacious concentration for a TCE is significantly more challenging than for a compound with a typical Emax efficacy curve.

One of the key steps during the early stages of drug development is lead compound selection, which is based on many criteria, including drug affinity for the target, denoted as an inverse of dissociation constant *K*_*D*_ and defined as 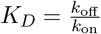, where *k*_on_ is the rate constant at which the drug reversibly binds to its target, and *k*_off_ is the dissociation rate constant. The question of affinity optimization to maximize the drug’s efficacy is particularly challenging for TCEs, even those targeting similar or even the same targets. For instance, REGN5458 and REGN5459 are both targeting CD3 and BCMA receptors but with different affinities for CD3 [30]. While tighter binding (lower *K*_*D*_) can be associated with achieving greater target engagement with lower drug concentration (higher potency), for TCEs and other drugs, whose efficacy is predicated on maximizing trimer concentration *Y*, this is not necessarily the case. Specifically, as was reported in [19, 31], very tight binding on each arm can “take up” most of the drug, leaving less drug available to bind to the other arm, thereby decreasing the probability of trimer complex formation. Therefore, there likely exists an interplay of “optimal affinities” for both arms of the molecule, resulting in different variations of the *bell-shaped* relationship between the drug concentration and trimer complex [19].

Four parameters are critical to characterize such a bell-shape curve at the site of action: 1) initial concentration of *T*_1_, 2) initial concentration of *T*_2_, 3) binding affinity/dissociation constant of the TCE to target 1 *K*_*D*1_, and 4) dissociation constant *K*_*D*2_. The initial concentration of *T*_1_ depends on the number of T cells in the body. Most of the existing TCEs bind to CD3 (see Table 1), and the average turnover parameters for this particular target are relatively well understood [32]. The initial concentration of *T*_2_ depends on the tumor and can vary dramatically. A review of existing CD3-based TCEs, summarized in Table 1, showed that the initial concentration of tumor-specific targets is generally considered to be less than 5 × 10^3^/cell; this however can vary across different cancer types.

The most desirable targets on cancer cells are minimally expressed in normal cells, so as to minimize off-target effects.

The affinity of the TCE molecule to *T*_1_ (CD3 on the T cells) can be significantly smaller, comparable, or significantly larger than the binding affinity to *T*_2_ (targeted receptor protein on the cancer cells). For example, the binding affinity of Blinatumomab [20], PF-06671008 [23], and 7370 [25] to *T*_1_ is lower than their affinity to *T*_2_. The affinity of Acapatamab [28], Solitomab [21], and PF-07062119 [26] to their targets are in a comparable range. On the other hand, TCE molecules such as BAY2010112 [22], Taralatamab [27], and REGN5458 [33] have a significantly higher affinity (lower dissociation constant) to *T*_1_.

The binding kinetics of the TCE molecules with respect to each of the targets may also affect the distribution of this type of antibody in different tissues. For instance, the biodistribution of CD3/HER2 TCE, also known as T-cell–Dependent Bispecific (TDB), has been measured for different ranges of affinities to CD3 for solid tumors in mouse models [24], confirming the intuition that higher affinities to CD3 (lower dissociation constant *K*_*D*1_) increase drug uptake in the peripheral tissues including lymph nodes, thereby decreasing their availability in the TME. Therefore, TCE molecules with lower affinities for CD3 are less toxic in treatments designed for solid tumors.

The characterization of the bell-shape was analytically investigated by [31]. The integration of the three-body model with the pharmacokinetics and efficacy of PF-06671008 TCE molecule was presented in [19]. Here, we build on these results to create a framework to allow quantitative comparison between different TCE molecules based on their bell-shape response. We anchor the analysis to the affinity values of several existing TCE molecules, and showcase the resulting bell-shape response and its sensitivity to different parameters. We further explore the identifiability of affinity values from the experimental data, and made the connection between pharmacokinetics, efficacy and biodistribution of the TCE based on an example, where TCE molecules that target HER2 were designed to have different affinities to CD3 receptor [24].

In what follows, the general properties of the two targets are assumed to be known, and the focus is on approaches to identify *K*_*D*_ values from preclinical experiments, and on characterizing their effect on the bell-shape pattern of the concentration trimer *Y* at the site of action. A quantitative comparison is presented for the TCE molecules that were available in the literature at the time of this writing. The computational analysis was performed for different scenarios, including varying ratios between target concentrations to quantify the sensitivity of projected efficacy (here correlated with maximizing the concentration of trimer *Y*) to initial conditions. The paper concludes with the formulation of a full quantitative pharmacology model that should facilitate a broader discussion of biodistribution of TCE molecules.

## 2 Methods

### 2.1 Three-body model

The initial process of lead compound selection is based on *in vitro* experiments, where cells are co-incubated with the drug in order to evaluate their relative affinities and where drug clearance does not affect the dynamics. Consequently, one can initially focus on just the nearly instantaneous formation of drug-target dimers and trimers, which can be interpreted as a three-body model. In recent investigations such as [19] it has been emphasized that the efficacy of the TCE drug trimer concentration at the site of action is a bell-shaped function. A too low or too high concentration of a TCE does not lead to formation of a sufficient number of trimer complexes: at low concentrations, too few trimers are formed, while at high concentrations the equilibrium shifts towards increased formation of dimers *D*_1_ and *D*_2_ and away from trimers. There exist, therefore, a “sweed spot” that maximizes trimer formation and consequently the expected efficacy.

A “three-body model” in the terminology of [31] is what all bispecific antibodies have in common. In the three-body model, a bispecific antibody (binding species) connects to two different target molecules (terminal species) to form a ternary complex. After the formation of dimers between the first target and the antibody, the binding kinetics can be changed based on a cooperativity factor to increase/decrease the binding affinity of the antibody molecule of the formed dimer to the second target. A positive cooperativity can be interpreted as avidity [34, 35, 36, 37], where the apparent affinity to the second target is greater once the drug has been bound to the first target, given that both targets are expressed on the same cell surface. The cooperativity factor can be neglected in models of bispecific antibodies that target receptors on two different cell types, such as TCEs.

As mentioned above, the model described in Figure 1 includes six state variables to describe the concentrations of the TCE *X*(*t*), the target receptor on the surface of the immune cells *T*_1_(*t*), the target receptor on the surface of cancer cells *T*_2_(*t*), the dimer complex of immune cell-antibody *D*_1_(*t*), the dimer complex of antibody-immune cell *D*_2_(*t*), and the ternary complex of immune cell-antibody-cancer cell *Y* (*t*). The initial concentration of the dimer and trimer complexes are assumed to be zero at the starting point *D*_1_(0) = *D*_2_(0) = *Y* (0) = 0. The initial concentration of the TCE *X*(0) is the concentration that is going to be assessed in subsequent analysis. The system of Ordinary Differential Equations (ODEs) describing the dynamics of these six variables is as follows:

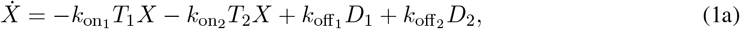

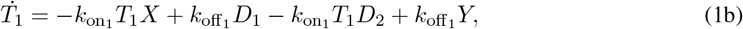

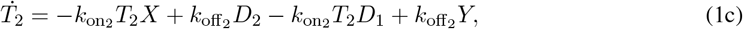

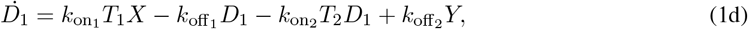

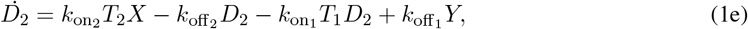

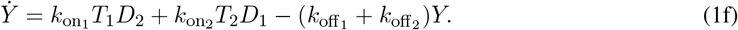

Note that the time dependence of the state variables is dropped for writing simplicity. The dot sign on top of each state variable on the left side represents the time derivative.

Model 1 contains four parameters: 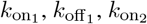 and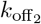. However, in preclinical measurements, using for example surface resonance experiments, one only estimates the dissociation constants of the two binding sites:

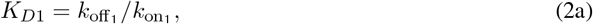

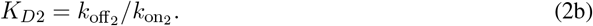

The dissociation constants *K*_*D*_ are sufficient to estimate the equilibrium concentration of each species in a dimerization reaction (for example 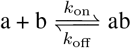), and different values of (*k*_on_, *k*_off_) that result in the same dissociation constant can change the reaction rate but do not affect the equilibrium concentrations.

The steady state reached when performing a long-time simulation of the three-body model (1) will predict the actual equilibrium concentrations, which can be thought of as an *in silico* equivalent of an *in vitro* dose-response experiment used to estimate projected target occupancy as a function of drug concentration. Figure 2 shows a numerical simulation of the system, with corresponding parameter values listed in Table 2, representing the PF-006671008 molecule [19], and a total simulation duration of 1000 hours. The y-axis in Figures 2a and 2b is normalized to capture % of Maximum Fluorescence Intensity (MFI), where MFI represents fluorescence intensity normalized to the maximum observed value in a given experiment. Because the normalization depends on the tested concentration range, MFI-based results may appear different across experiments even if the underlying biological response is similar. The relative concentrations are measured based on the relative strength of the fluorescent signal. Analyzing the factors that affect the characteristic bell shape of the projected trimer concentration is the focus of this section. Picking the “right” range of molecule concentrations is critical in visualization of the bell shape. For example, note when comparing Figure 2a and Figure 2b, how different experimental results might look when reported in MFI format with different ranges of drug concentration on the x-axis.

**Table 2:** Parameters used in model (3), are based on previously published parameters [19]; parameters different from [19] are marked by asterisk.

|  | Value | Unit | Definition |
| --- | --- | --- | --- |
| $V_c$ | 40.2 | mL/kg | Volume of distribution in the central compartment |
| $V_p$ | 211 | mL/kg | Volume of distribution in the peripheral compartment |
| $V_l^*$ | 92 | mL/kg | Volume of distribution in the lymph compartment |
| $k_{elX}$ | $1.16 \times 10^{-1}$ | 1/h | Clearance in the central compartment |
| $k_{elT}$ | 2.51 | 1/day | Elimination rate of immune cells (Target 1) |
| $k_{cp}$ | $6.27 \times 10^{-1}$ | 1/h | Redistribution rate constant to the peripheral compartment |
| $k_{pc}$ | $1.19 \times 10^{-1}$ | 1/h | Redistribution rate constant to the central compartment |
| $k_{clT}^*$ | $1 \times 10^{-1}$ | 1/day | Target 1 redistribution to the lymph nodes |
| $\alpha$ | 1 | 1/nM | Empirical parameter |
| $\beta$ | 5 | 1 | Empirical parameter |
| $k_{lcT}^*$ | $5 \times 10^{-1}$ | 1/day | Target 1 redistribution to the central from lymph |
| $k_{cl}^*$ | $5 \times 10^{-2}$ | 1/day | Redistribution rate constant to the lymph compartment |
| $k_{on1}$ | 1.72 | 1/nM/h | Binding of the antibody and target 1 |
| $k_{off1}$ | 19.66 | 1/h | Unbinding rate of the antibody-target 2 dimer |
| $k_{on2}$ | 1.57 | 1/nM/h | Binding rate of the antibody and target 2 |
| $k_{off2}$ | 0.74 | 1/h | Unbinding rate of the antibody-target 2 dimer |
| $k_{deg}$ | $1.5 \times 10^{-1}$ | 1/h | Tumor $T2_c$ degradation rate constant |
| $k_{syn}$ | $k_{deg} \times T2_c(0)$ | nM/h | Tumor synthesis rate constant |
| $k_{synT}$ | $k_{elT} \times \frac{T1_c(0)V_l}{V_c}$ | nM/h | Synthesis rate of the immune cells (no proliferation) |
| $k_{td}$ | 1.082 | 1/h | Disposition rate to the TME (depends on tumor physiology [38]) |
| $k_{ctT}$ | $2 \times 10^{-3}$ | 1/day | T cell redistribution from the central to the TME |
| $k_{tcT}$ | $5 \times 10^{-4}$ | 1/day | T cell redistribution from the TME. |
| $k_{max}$ | 1.32 | 1/day | Maximum killing rate |
| $k_{c50}$ | $6.9 \times 10^{-5}$ | nM | Concentration at half maximum |
| $k_{ge}$ | $1.9 \times 10^{-1}$ | 1/day | Exponential tumor growth rate |
| $k_{gl}$ | $1.23 \times 10^{-1}$ | mL/day | Linear tumor growth rate |
| $k_v$ | 6.0 | mL | Maximum tumor volume |
| $k_\tau$ | 3.99 | day | Transduction time between tumor compartments. |
| $k_\psi$ | 20 | 1 | Exponential to linear transition rate of the tumor |
| $w$ | 80 | kg | Weight of a patient. |
| $\rho$ | 9.52 | nM/(μg/kg) | Concentration unit conversion for in the central compartment |

**Figure 2:**
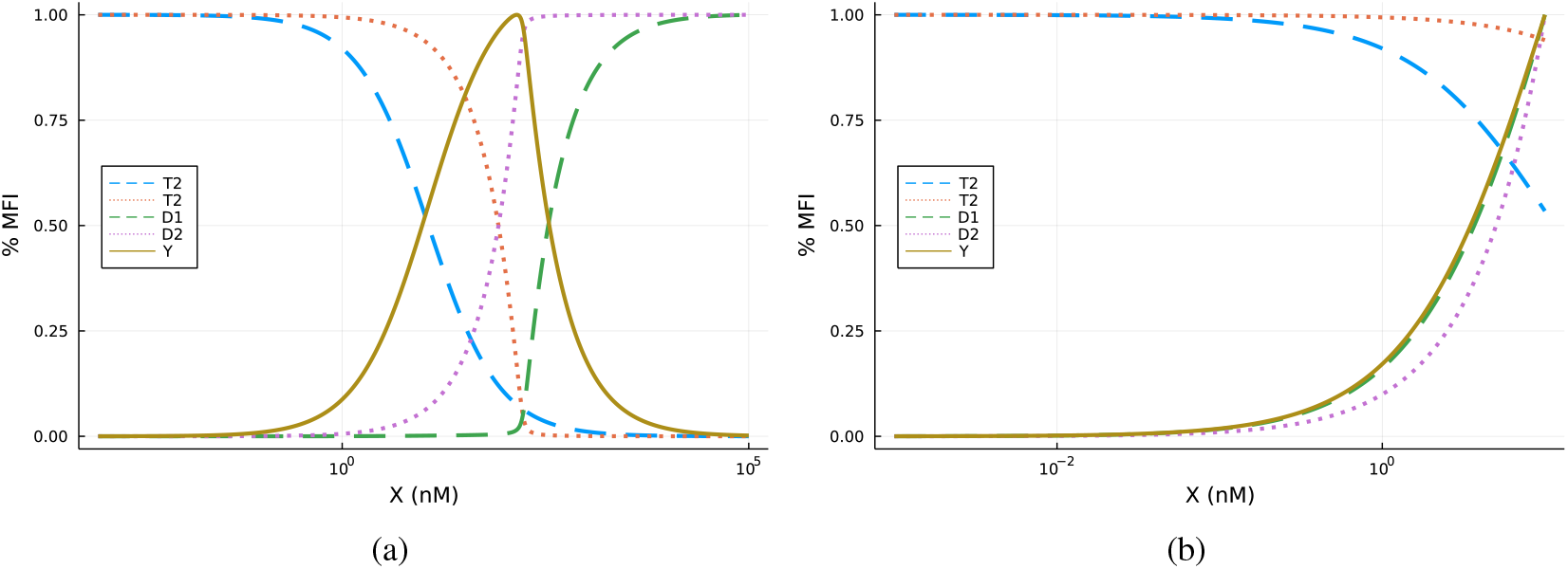
Model-based steady state simulations of the three-body model (1). The vertical axis is normalized to % MFI, which represents relative fluorescence intensity normalized to the maximum observed value in a hypothetical experiment. This figure does not represent experimental data, but rather simulations using the model under two different TCE concentration ranges. These simulations illustrate that if one were to conduct experiments using different concentration ranges, they might observe distinctive patterns such as those shown here, even if the underlying biological processes are consistent. The variables in the simulations have different ranges of initial concentration of the TCE. The initial TCE concentration range and the range of horizontal axis values in (a) are larger than in (b). The plots on the left show the concentrations of the trimer and the targets, and the plots on the right side show the drug-target dimer concentrations.

### 2.2 The bell-shaped response

We have investigated the sensitivity of the bell-shaped response peak, to the binding kinetics of the TCE molecule at the site of action, here in the TME, and the initial concentration of each target. The maximum concentration (peak) of the trimer complex *Y* and the corresponding initial TCE concentration that results in maximum trimer concentration (optimal TCE) are the main outputs of interest in this analysis. We used the three-body portion of the quantitative model published for PF-06671008 TCE molecule [19], as an example, to assess the sensitivity of the peak of the bell-shape by changing each of the parameters in model (1), while keeping the other parameters of the model constant.

### 2.3 Comparison between TCEs

The three-body model (1) was used as a tool to quantitatively compare existing CD3-based TCEs summarized in Table 1 based on their published binding kinetics to their targets. The molecules included in this study are divided into two general categories: targeting solid tumors and targeting liquid tumors. Although each of the molecules might have different structures, biodistribution, metabolism, or pharmacokinetic characteristics, the three-body model captures the general structure of the interactions of these compounds with their targets at the site of action.

Numerical simulations were conducted assuming initial target concentrations of 0.108 nM and 166 nM (based on [19]). Due to the lack of reported association (*k*_on_) and dissociation (*k*_off_) rate constants for all molecules, *k*_on_ was fixed to 1 nM^−1^ h^−1^, and *k*_off_ was inferred from the published equilibrium dissociation constant (*K*_*D*_ × 1 nM^−1^ h^−1^) values listed in Table 1.

### 2.4 Biodistribution

A full dynamic model for biodistribution of the TCE molecules in humans with IV dosing is shown in Equations (3). This is a modified version of the model presented in [19], where the number of tumor-related compartments is reduced to two for simplicity, and all the immune system tissues such as lymph nodes are lumped into a single additional compartment. We use this model to obtain a better mechanistic understanding of the relationship between dissociation kinetics and biodistribution.

The differences between model (3) and the model presented in [19] are as follows: 1) an extra compartment is included in order to represent tissues with a higher concentration of immune cells, e.g. lymph nodes, 2) intermediate compartments are integrated into the main tumor compartment, and 3) biodistribution depends on the affinity of the TCE molecule to different targets, which could be explained by the flow of the immune cells between lymph nodes and the central compartment. The additional parameters included in this model are not fit to data, and the presented parameters are chosen arbitrarily based on numerical simulations to demonstrate the quantitative approach for explaining the biodistribution of the TCE molecules. Molecule and target-specific parameters can be used as needed provided data availability.

The relative change of the target concentration is assumed to be negligible in comparison with the distribution rate constants of the TCE in the present model. Moreover, the rate constant of target 1 is assumed to be faster between the central compartment and the lymph nodes (*k*_*lcT*_ and *k*_*clT*_) relative to the rate constant of the target 1 between the central compartment and the TME (*k*_*ctT*_ and *k*_*tcT*_). The rate constants of the TCE between the central compartment and the lymph nodes (*k*_*lc*_ and *k*_*cl*_) are assumed to be slower than the rate constants of target 1 (*k*_*lcT*_ and *k*_*clT*_). The reason is the increased flow rate of immune cells between lymph nodes and the central compartment.

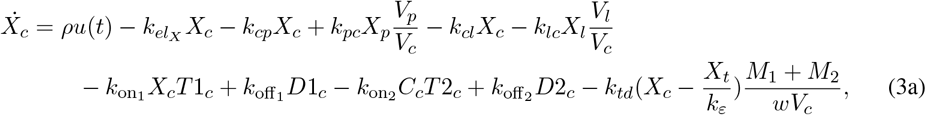

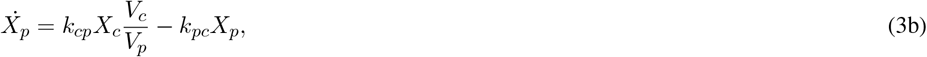

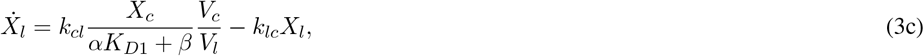

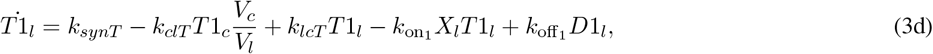

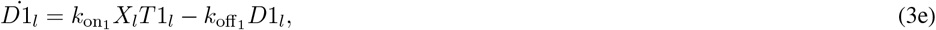

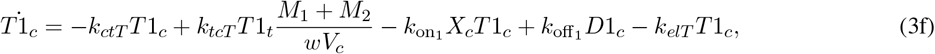

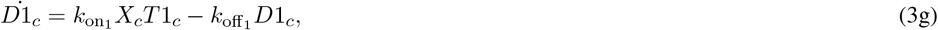

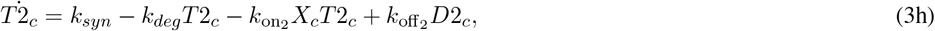

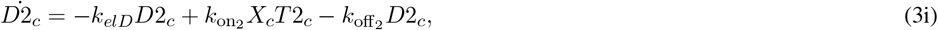

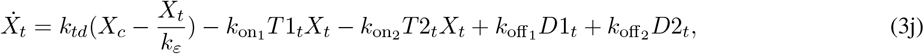

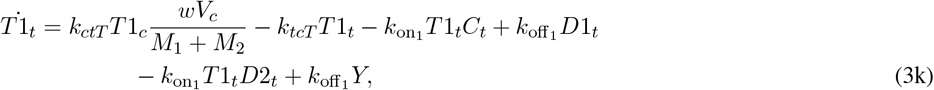

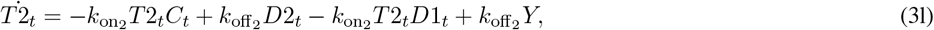

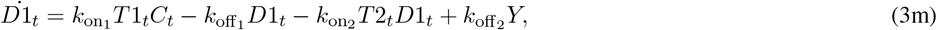

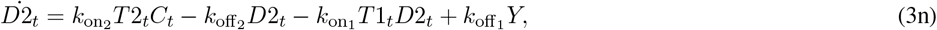

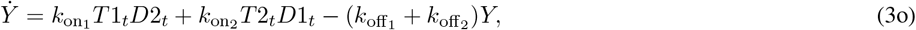

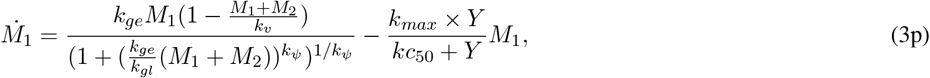

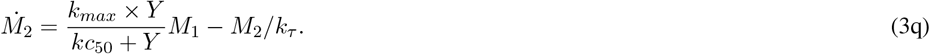

In these equations, *X* is the concentration of drug, *T* is the concentration of targets, *D* is the drug-target dimer concentration, *Y* is the trimer concentration, and *M* is the tumor volume intermediate compartment. All parameters and variables are defined in detail in Tables 2 and 3. The dot sign on top of the variables is a time derivative 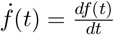.

**Table 3:**
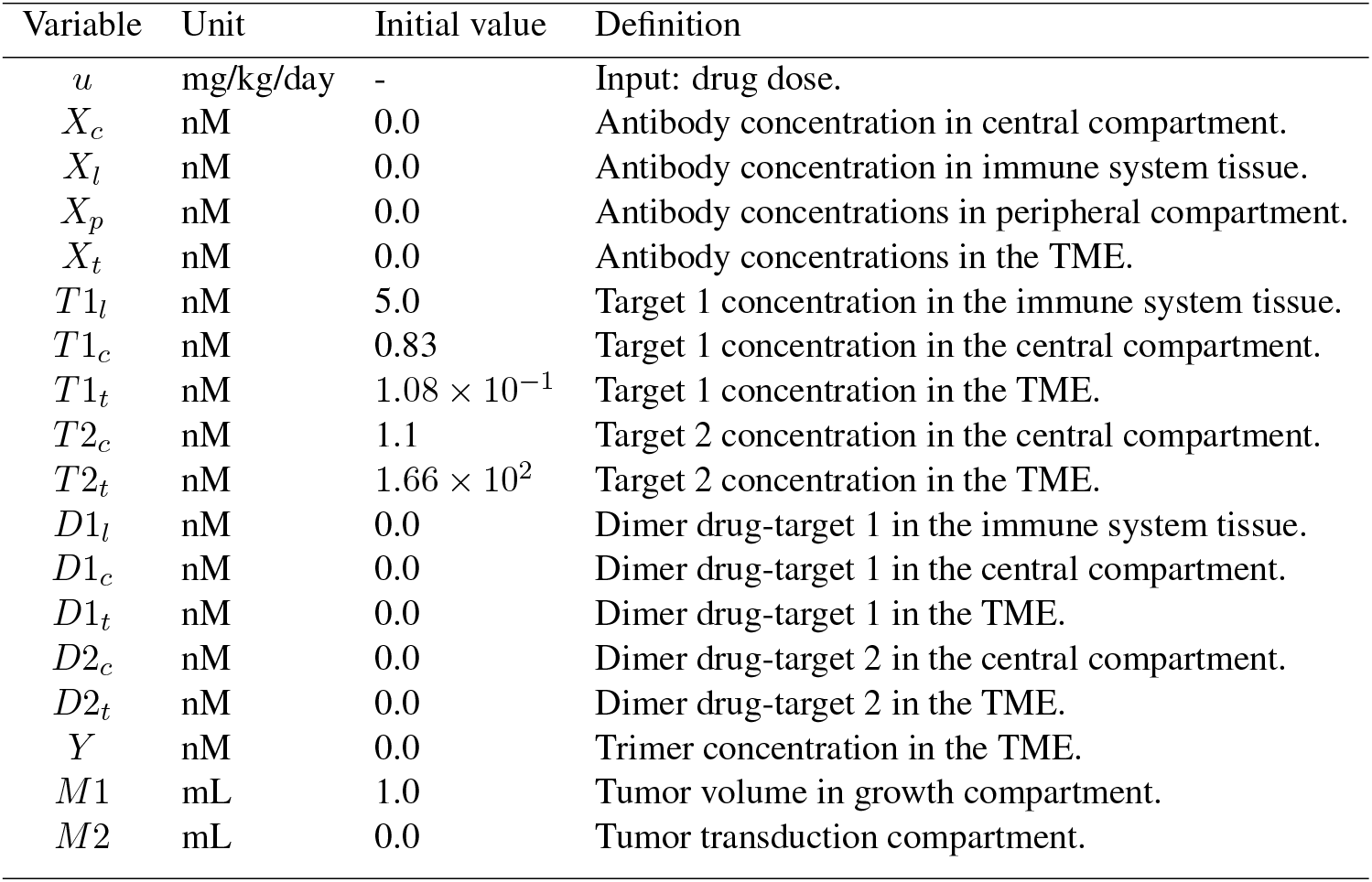
Variables used in model (3).

| Variable | Unit | Initial value | Definition |
| --- | --- | --- | --- |
| $u$ | mg/kg/day | - | Input: drug dose. |
| $X_c$ | nM | 0.0 | Antibody concentration in central compartment. |
| $X_l$ | nM | 0.0 | Antibody concentration in immune system tissue. |
| $X_p$ | nM | 0.0 | Antibody concentrations in peripheral compartment. |
| $X_t$ | nM | 0.0 | Antibody concentrations in the TME. |
| $T1_l$ | nM | 5.0 | Target 1 concentration in the immune system tissue. |
| $T1_c$ | nM | 0.83 | Target 1 concentration in the central compartment. |
| $T1_t$ | nM | $1.08 \times 10^{-1}$ | Target 1 concentration in the TME. |
| $T2_c$ | nM | 1.1 | Target 2 concentration in the central compartment. |
| $T2_t$ | nM | $1.66 \times 10^2$ | Target 2 concentration in the TME. |
| $D1_l$ | nM | 0.0 | Dimer drug-target 1 in the immune system tissue. |
| $D1_c$ | nM | 0.0 | Dimer drug-target 1 in the central compartment. |
| $D1_t$ | nM | 0.0 | Dimer drug-target 1 in the TME. |
| $D2_c$ | nM | 0.0 | Dimer drug-target 2 in the central compartment. |
| $D2_t$ | nM | 0.0 | Dimer drug-target 2 in the TME. |
| $Y$ | nM | 0.0 | Trimer concentration in the TME. |
| $M1$ | mL | 1.0 | Tumor volume in growth compartment. |
| $M2$ | mL | 0.0 | Tumor transduction compartment. |

### 2.5 Identifiability

The goal of performing identifiability analysis is to assess whether model parameters could be uniquely recovered from experiments. In this paper we will discuss two different setups for identifiability: 1) identification of the *K*_*D*_ values from time course data, and 2) identification of *K*_*D*_ values from steady state data. Time course data plays a major role in the drug development process for *in vivo* studies, and steady state data is often used for *in vitro* studies to assess potential drug efficacy. Notably, this analysis is based on deterministic data and should be treated as a proof-of-concept approach to recover *K*_*D*_ values from specific experiments.

The standard context for performing identifiability analysis is for the case when the parameter values are to be inferred from time course data for some measured quantities [39]. In this case, one typically considers an ODE model in the state-space form

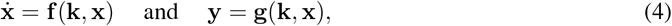

where **x** and **y** are the vectors of states and observed quantities, respectively, and **k** is a vector of scalar parameters. For the three-body model (1), we would have **x** = [*X, T*_1_, *T*_2_, *D*_1_, *D*_2_, *Y*]^*T*^, the observed quantities would be *y*_1_ = *X, y*_2_ = *T*_1_, *y*_3_ = *T*_2_ (free drug, first target, and second target, respectively), and 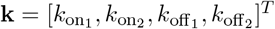.

For a model (4), a function *h*(**k**) of parameters (could be just a parameter) can be

- *globally identifiable* if, for almost every solution of (4), any other solution of (4) with the same trajectory for **y** must have the same value of *h*(**k**).
- *locally identifiable* if, for almost every solution of (4), there are finitely many possible values of *h*(**k**) for any other solution of (4) with the same trajectory for **y**;
- *nonidentifiable* if, for almost every solution of (4), there are infinitely many possible values of *h*(**k**) for other solutions of (4) with the same trajectory for **y**.

Let us elaborate on the notion of “almost every” in the above definitions. Every trajectory of (4) is uniquely determined by a pair (**k, x**(0)) of the parameter vectors and initial conditions. Then “almost every” means that that there exists a manifold of codimension at least one (and, thus, of Lebesque measure zero) in this (**k, x**(0))-space such that the stated property may fail only for the trajectories from this manifold. For a more precise formalization of this notion and related discussions, we refer to [40, Section 2]. Note that a globally identifiable parameter is locally identifiable as well. The term “*generic*” (global or local) identifiability is sometimes used in order to emphasize that there is a possible exceptional set in which identifiability does not hold. We will give two simple examples to illustrate these notions.

#### Example 1

Consider a model 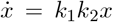 with the observable *y* = *x*. Then the trajectories are of the form 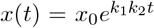. We can see that neither *k*_1_ nor *k*_2_ is identifiable because the same trajectory can be produced by infinitely many triples (*k*_1_*/c, k*_2_*c, x*(0)), where *c* ranges over nonzero numbers. On the other hand, take the function *h*(**k**) = *k*_1_*k*_2_. Consider the manifold consisting of all pairs of the form (**k**, 0), that is, pairs of parameter vectors and the special initial state *x*(0) = 0. For every solution not starting from this manifold, we have identifiability. Indeed, if 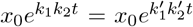 for all *t*, and *x*_0_ ≠ 0, then 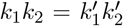. Thus, *h*(**k**) is globally identifiable.

#### Example 2

Consider a model 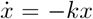 with the observable *y* = *x*. Then the trajectories are of the form *x*(*t*) = *x*_0_*e*^−*kt*^. If *x*(0) ≠ 0, the growth rate *k* is uniquely defined by the function *y*(*t*) via, for example, a formula 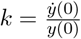. If *x*(0) = 0, then all values of *k* yield the same *y*(*t*) = 0.

We claim that in this model *k* is globally identifiable. Indeed, the trajectories not allowing for unique reconstruction of *k* (namely, the trajectories with *x*(*t*) = 0) correspond to a codimension one subvariety (*k*, 0) in the (*k, x*(0))-space parametrizing all the trajectories.

There are a number of software tools for assessing identifiability [41] using the time course data. We have chosen to use the web-based Structural Identifiability Analyzer [42], which is based on the algorithms from [43, 44]. This algorithm takes as input a model in the format (4)^1^. For the three-body model (1), the software showed that all four parameters 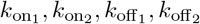 are globally identifiable from the time course data.

While the time course identifiability results presented above are of mathematical interest, they are not the most useful in our context, because most experimental measurements are typically done at steady state [20, 21, 22, 23, 25, 26, 27, 28]. Thus, we next turn our attention to the more interesting question of identification on the basis of steady state data alone.

In this setup, we fix a subset 𝒯 in the (**k, x**(0))-space of the considered trajectories (e.g., trajectories with positive parameter values and nonnegative initial conditions). For the problem to be well-posed, it is necessary that every trajectory in 𝒯 converges to a steady state. For (**k, x**(0)) ∈ 𝒯, we will denote the corresponding steady state by 𝒮(**k, x**(0)). Then we will call a function *h*(**k**) globally identifiable from the steady state data if there exists a subset 𝒯_0_ ⊂ 𝒯 of Lebesque measure zero such that

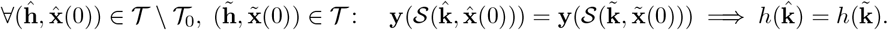

We perform a detailed analysis of the three-body model in this setup in Section 3.4, and here we give illustrative examples explaining the steady state setting and its differences from the time course identifiability.

#### Example 3

We revisit Example 2 with the model 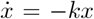, and set 𝒯 to be trajectories with positive *k*. Then, for any initial condition, there is a unique attracting steady state 𝒮 (*k, x*(0)) = 0. Assume that what is measured is precisely this steady state. Since the steady state is zero regardless of the value of *k*, the parameter *k* is not identifiable from the steady state data (unlike the time course setting of Example 2).

#### Example 4

Consider the model of logistic growth 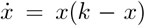 with *y* = *x* and 𝒯 being the set of (*k, x*(0)) with *k >* 0 and *x*(0) ⩾ 0. It is well-known that, if *x*(0) *>* 0, there is a unique attracting steady state 𝒮 (*k, x*(0)) = *k*. Furthermore, we have 𝒮 (*k*, 0) = 0. Therefore, the knowledge of the steady state is sufficient to find the value of *k* if *x*(0) *>* 0. Thus, by setting 𝒯_0_ = {(*k, x*(0)) ∈ 𝒯 | *x*(0) = 0}, we see that *k* is globally identifiable.

The next example can serve as a toy model for the analysis of the three-body problem we will perform in Section 3.4.

#### Example 5

Consider the following chemical reaction network with three species *X*_1_, *X*_2_, *X*_3_:

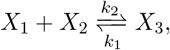

which, under the law of mass-action kinetics, is governed by the following ODE system (*x*_*i*_ denotes the concentration of *X*_*i*_ for *i* = 1, 2, 3):

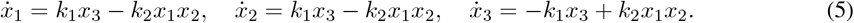

We will consider the set 𝒯 of trajectories with positive parameter values and *x*_1_(0) = *c*_1_ *>* 0, *x*_2_(0) = *c*_2_ *>* 0, *x*_3_(0) = 0. It is known [45] that every such trajectory converges to a steady state. We assume that the measured quantities are the values of *x*_2_ and *x*_3_ at the steady state, that is, **y** = [*x*_2_, *x*_3_]^*T*^, and we are interested in identifiability of 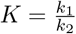.

If 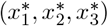 are the coordinates of the steady state, then they satisfy the equation

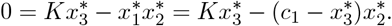

The equation contains two unknown values, *K* and *c*_1_, so we cannot find the value of *K* using only this information.

Assume now that we can perform two independent experiments with the same (but still unknown) value of *c*_1_ and different values of *c*_2_, we will denote them 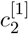 and 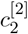. Formally, we have model consisting of two copies of (5) and the set of trajectories

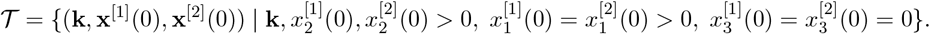

We denote the coordinates of the steady states by 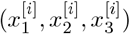 for *i* = 1, 2. In this case we get a system of two equations:

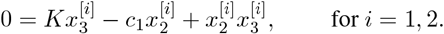

Regarding this system as a linear system in the unknowns *c*_1_ and *K*, we can use Cramer’s rule to eliminate *c*_1_ and get a formula for *K* (this elimination task becomes more tedious in higher dimensions, so we will employ Gröbner bases to do this in Section 3.4):

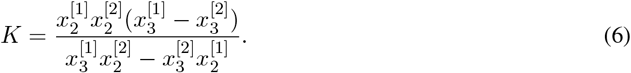

The formula above is well-defined as long as 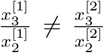. The value of 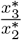 is completely determined by *K, c*_1_, *c*_2_. Furthermore, it is an algebraic function in these variables (see [45, p. 1036]). We observe that this function is nonconstant with respect to *c*_2_ when *c*_1_ and *K* are fixed. Indeed, since 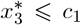 and 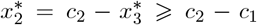, the value of the function can be made arbitrarily close to zero by taking *c*_2_ sufficiently large. Therefore, for every 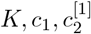, there are only finitely many values of 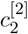 with 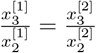.

Therefore, a subset 𝒯_0_ of pairs trajectories for which 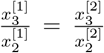 is of measure zero inside 𝒯. Together with (6), this proves identifiability of *K*. So we conclude that the value of *K* is identifiable from the steady state data of two experiments.

### 2.6 Computational resources

SIAN [43] was used for structural identifiability analysis and analytical derivations. Numerical simulations and figures are produced with the Julia programming language [46]. The DifferentialEquations package was used for numerical calculations [47], with Tsit5() numerical algorithm that is suitable for non-stiff problems. The numerical software for reproducing the figures presented in the manuscript along with more examples are publicly available on Github [48].

## 3 Results

### 3.1 Results regarding the bell-shaped response

We examined the bell-shape response using the molecule PF-06671008 TCE from [19] as an example to assess the sensitivity of the peak of the bell-shape to the *K*_*D*_ values between the drug and either of its two targets (see Figure 3). As one can see in the two left subfigures of Figure 3a an increase in the values of *K*_*D*1_, and *K*_*D*2_ results in a significant decrease of the maximal value of *Y*. On the other hand, by looking at the right side of Figure 3a, it can be observed that the optimal concentration of TCE (the corresponding initial concentration of the TCE at the peak of the trimer concentration) will be decreased by only reducing *K*_*D*1_, and increased by increasing either *K*_*D*1_ or *K*_*D*2_.

**Figure 3:**
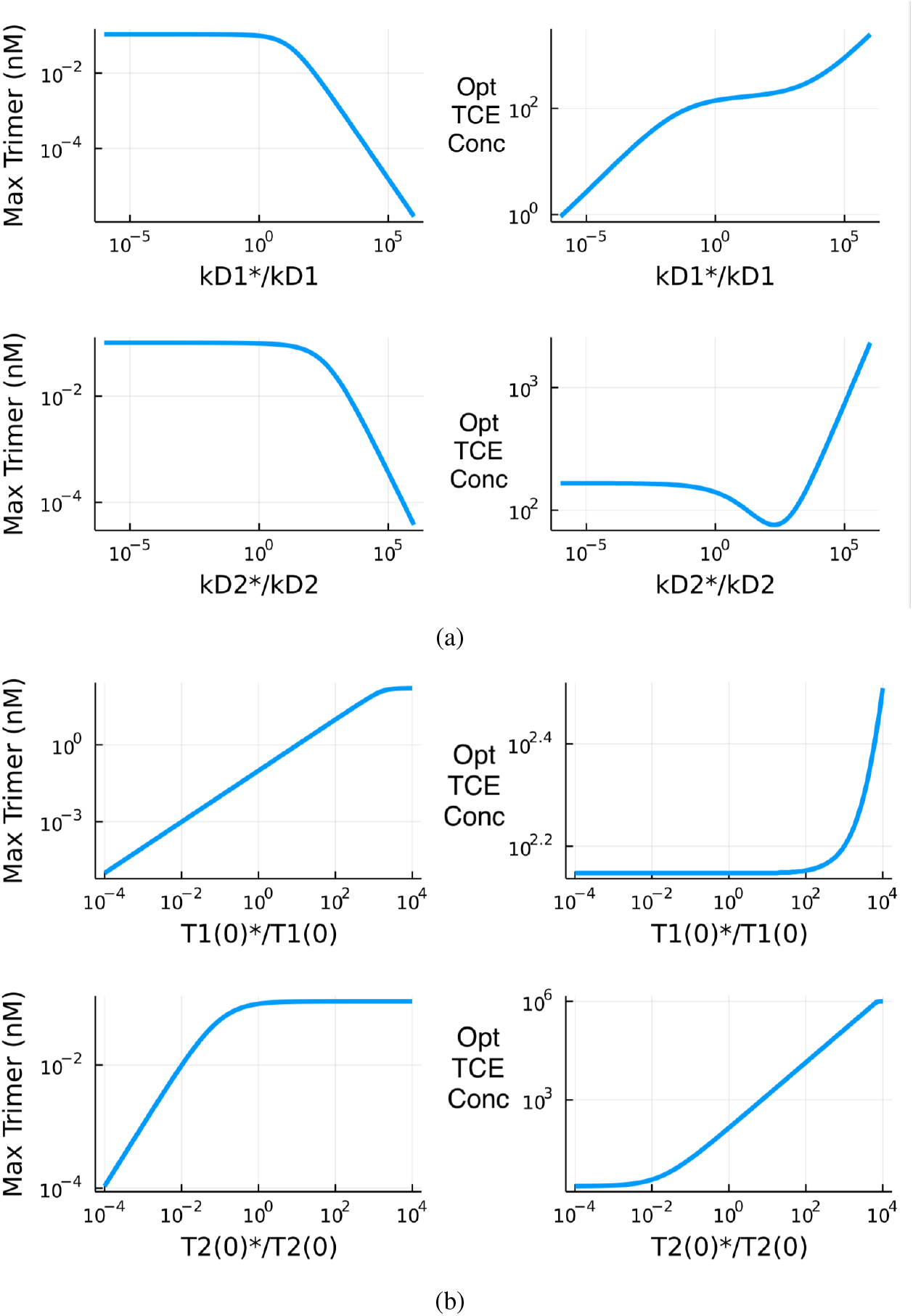
Log-log plots of the bell-shape characteristics: the effects of (a) sweeping dissociation constants, and (b) sweeping target concentrations at the site of action on the peak of the trimer concentration *Y* and it corresponding TCE concentration *X*. The horizontal axes are the log scale difference between the modified parameter (marked with a star*) and their original value. Maximum trimer concentration is the peak of the bell-shape, and Optimal TCE is the corresponding initial antibody concentration, based on model (1).

The sensitivity of the maximum trimer concentration, and optimal TCE initial concentration to the initial concentration of the targets on the immune cells *T*_1_(0), and cancer cells *T*_2_(0) are visualized in one-dimensional plots of Figure 3b. From the left side, it can be observed that the maximum trimer concentration is sensitive to *T*_1_(0) and insensitive to *T*_2_(0), which is physiologically reasonable, since the initial concentration of the first target, CD3 receptors on immune cells, is much smaller relative to the second target, P-cad protein on the tumor cells. This result is consistent with the sensitivity analysis presented in [19]. Surprisingly, any change above 0.01x, or below 100x in the initial concentration of the target *T*_1_(0) does not affect the predicted optimal TCE concentration but significantly changes the peak of trimer concentration *Y*.

The dissociation constants *K*_*D*1_ and *K*_*D*2_ depend on the TCE design, while *T*_1_(0) and *T*_2_(0) depend on the tumor characteristics and variability among cancer patients. From a design perspective, it would be ideal if the maximum trimer concentration and the corresponding TCE concentration are less sensitive to the initial concentration of the targets. From a toxicity perspective, it would be favorable if a lower concentration of TCE antibody can produce the same amount of trimer in the TME. In the analysis presented in Figure 3a, it can be observed that the concentration of TCE that results in the maximum trimer concentration can be decreased by reducing the dissociation constant of the first target *K*_*D*1_, and also that lower *K*_*D*1_ will result in a slightly higher trimer concentration.

Figure 4 extends the visualizations presented in Figure 3. The sensitivity of the maximum trimer concentration is on the left, and the sensitivity of the corresponding initial concentration of the TCE molecule is on the right. The red color represents a higher value of the nM concentrations in the log scale, and the blue color represents lower concentrations in the log scale. The vertical pattern in Figure 4a, and the horizontal pattern in Figure 4d are consistent with conclusions made from Figure 3. Moreover, the contrasting colors in the top left and the bottom right of Figure 4b suggest that a simultaneous increase in dissociation constants *K*_*D*1_, with a decrease in *K*_*D*2_, is favorable in reducing the required TCE concentration to achieve the maximum concentration of the trimer at the site of action.

**Figure 4:**
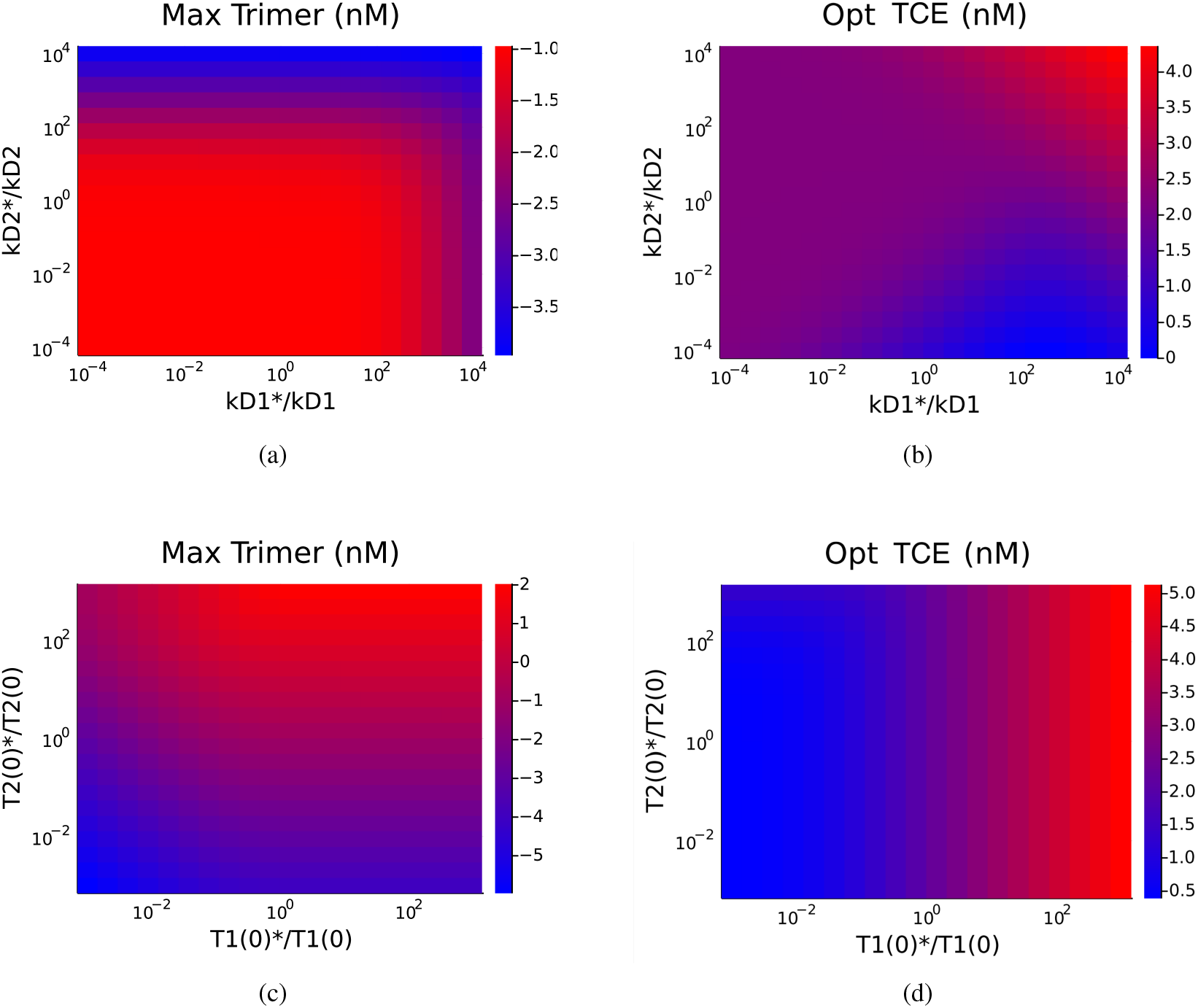
Bell-shape characteristics heatmap: the effects of sweeping the TCE antibody dissociation constants, *K*_*D*1_ and *K*_*D*2_, on (a) maximum of the trimer concentration of the bell-shape, and (b) the corresponding optimal, TCE concentration. The effect of sweeping initial target concentrations on (c) maximum of the trimer concentration of the bell-shape, and (d) the corresponding optimal concentration of the drug. The colors are plotted in log scale concentrations in nM. The horizontal and vertical axes are in log scale difference of the modified parameter (marked with a star*) and its original value, based on model (1).

### 3.2 Results on comparison between TCEs

We used the three-body model (1) in order to quantitatively compare the existing CD3-based TCEs summarized in Table 1. We grouped these molecules into two general categories: those targeting solid tumors and those targeting liquid tumors. In addition to *K*_*D*_ values summarized in Table 1, the initial concentration of the targets is necessary for a computational study of the three-body model. As the initial concentration of target receptors is highly dependent on cancer type, the comparison between the main characteristics of the bell-shape pattern, like the trimer peak, the corresponding TCE concentration at the peak, and the width of the peak, are done for different initial concentrations of the targets to capture a variety of scenarios.

A visual comparison between the bell-shapes of the molecules considered in this study is shown in Figure 5. The ratio of the initial concentration of the targets (*T*_1_(0) : *T*_2_(0)) is expected to be in the range of 1:10 to 1:1000 for solid tumors (Figure 5a), and in the range of 10:1 to 1:10 for liquid tumors (Figure 5b). It can be observed that in addition to the peak of the bell-shape, and the corresponding concentration of the TCE, the width of the bell-shape varies for different ratios of the initial concentration of the targets. For instance, the width of the peak of Tarlatamab is higher for a dense tumor, where the initial concentration of the target receptors on the cancer cells is greater than the concentration of target receptors on the immune cells.

**Figure 5:**
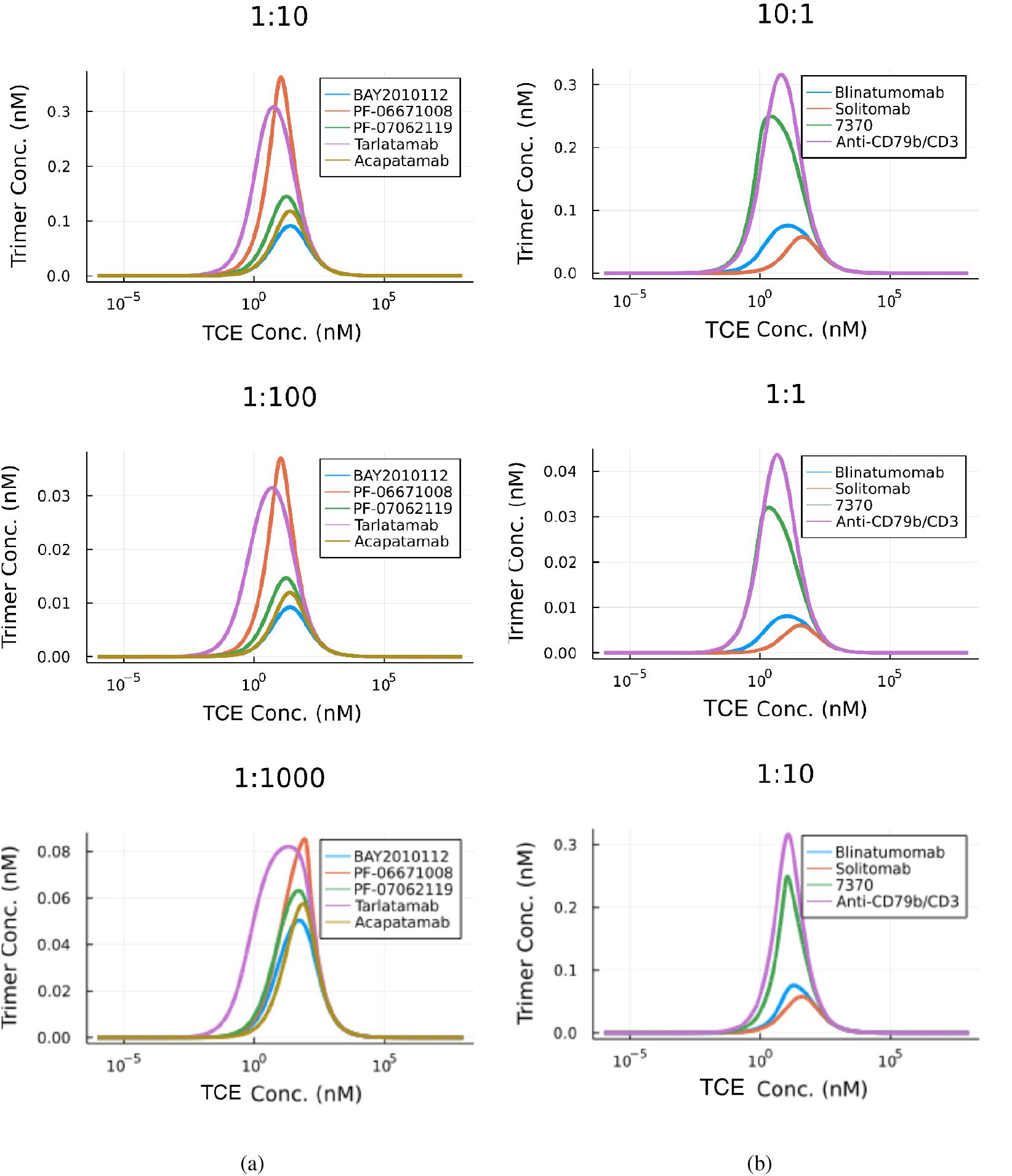
Numerical comparison between the bell shapes of TCEs designed for (a) solid tumors, and (b) liquid tumors based on model (1). The relative concentration of the targets *T*_1_(0) : *T*_2_(0) is printed on the top of each plot. The horizontal axes are in log scale, and the vertical axes are in linear scale.

The quantitative framework used for comparing bell-shapes of the TCE molecules at different ratios of initial concentration of the targets can be extended to continuous ratios of the targets. For this purpose, the basic characteristics of bell-shape are extracted across different ratios between the targets, shown in Figure 6. The peak of the trimer concentration is the maximum of bell-shape (on top), the corresponding TCE concentration is the initial concentration of the TCE that results in the maximum of the bell-shape (in the middle), and the bell-shape width is simply the range of the TCE concentration that results in at least 50% of the maximum of the bell-shape (on the bottom).

**Figure 6:**
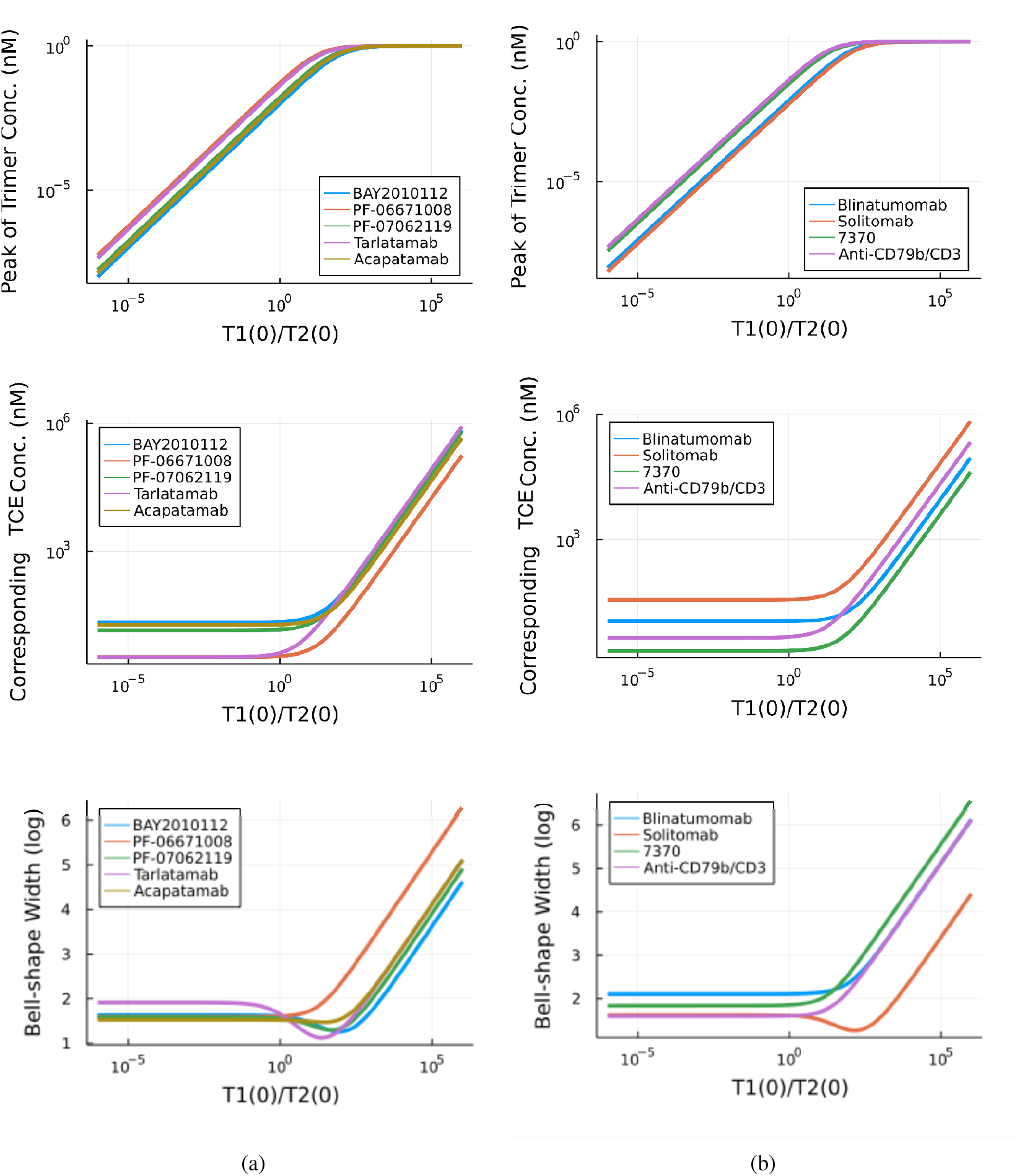
Numerical comparison between the basic characteristics of the bell-shape response of different TCEs designed for (a) solid, and (b) liquid tumors based on model (1). The top figures represent the value of the peak of bell-shape, the middle figures represent the corresponding TCE antibody concentration at the peak of the bell-shape, and the bottom figures represent the width of the peak of the bell-shape. The relative initial concentration of the targets *T*_1_(0)*/T*_2_(0) is visualized in log scale of the horizontal axes.

For a realistic comparison between TCEs for solid tumors (Figure 6a), the left-hand side of the horizontal axis should be considered, where the initial concentration of target receptors on cancer cells is much greater than the initial concentration of the CD3 target receptors on immune T cells. Similarly, for a realistic comparison between TCEs designed for liquid tumors (Figure 6b), the right middle or right side of the horizontal should be taken into consideration.

### 3.3 Results on biodistribution

It is clear that the biodistribution of TCEs is highly dependent on the parameters given in Table 3. Note that the parameters presented here are for a nominal TCE molecule, and are not identified from a specific data set.

In addition to dissociation constants, parameters affecting drug distribution can have a critical impact on trimer maximization. These parameters are essentially the disposition rate constant of the TCE to the lymph nodes *k*_*cl*_, and the disposition rate constant of the TCE to the TME. Notably, other parameters, such as the elimination rate 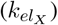 in the model, could also be affected by TCE design. Efforts to extend the TCE half-life led to the discovery of HLE-TCE molecules like Tarlatamab [27], and Acapatamab [28].

From the experimental results presented in [24], it can be observed that an increase in the affinity of the TCE toward immune cells (CD3, or target 1) increases its concentration in immune system-related tissues like lymph nodes. The mechanism of action suggests an inverse relationship between parameters *k*_*cl*_ and *K*_*D*1_. The numerical simulation presented in Figure 7a compares two nominal TCEs with different *K*_*D*1_ values administered at the same dose level. While pharmacokinetics in the central compartment *X*_*c*_, and the peripheral compartment *X*_*p*_ are similar, the concentrations at the lymph compartment *X*_*l*_ are different. Furthermore, different affinities of the TCE molecule to different targets lead to a different trimer concentration in the TME, which is discussed in detail in the previous section. Model 2, with parameters and initial conditions specified in Tables 2 and 3, was used to simulate these two molecules. The only difference between the two simulations is the value of 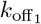, which was adjusted to reflect a 10-fold difference in *K*_*D*1_.

**Figure 7:**
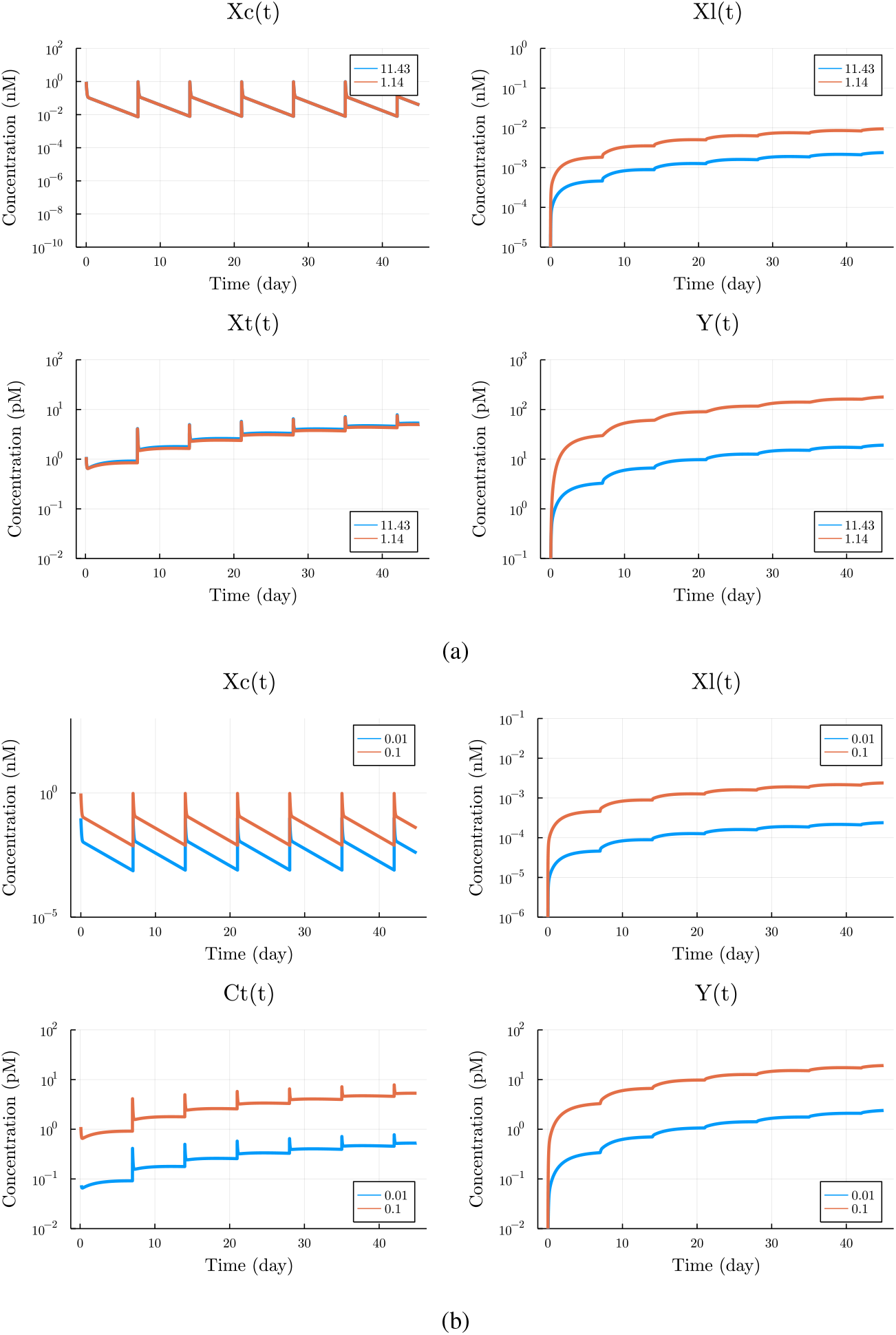
Numerical simulations of the biodistribution model (3) for (a) different dissociation constant *K*_*D*1_ between TCE and immune cells and the same dose (labels represent *K*_*D*1_ in nM), and (b) different dose of TCE and the same dissociation constant (labels represent dosing levels in mg/kg).

To better understand how a ten-fold change in the affinity can be compared with a ten-fold change in the drug dose, Figure 7b is presented. Notably, parameters *α* and *β*, defined as empirical parameters related to the lymph compartment, update the relationship with the dissociation constant *K*_*D*1_.

### 3.4 Results on identifiability from steady state data

In this section, it is proved that the dissociation constants 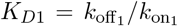and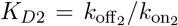 are identifiable from just three generic steady state measurements that are different in the initial concentration of the TCE molecule with equal initial concentrations of the targets. By generic we mean that, for each exeperiment, there are only finitely many values of the initial concetrations to avoid; for the precise statement, see Theorem 1.

Several approaches to analyzing identifiability from steady states have been developed, see [49, 50], but they are not applicable directly to our specific setup (due to limited observations, multiple experiments with shared conserved quantities, etc).

Some of the proofs in this section rely on tedious symbolic computations which were performed on a computer. The corresponding code can be found in the *identifiability* folder of the repository [48] with the supplementary code for the paper.

In terms of state variables, consider a steady state 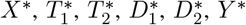 of the system (1). Then the numbers 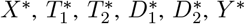, and 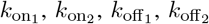 are related by a system of six polynomial equations obtained by setting the left-hand sides of (1) to zero, that is:

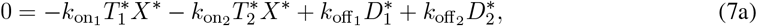

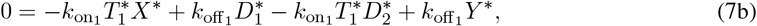

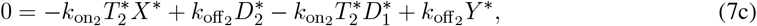

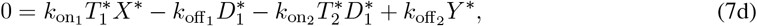

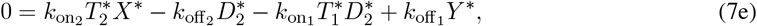

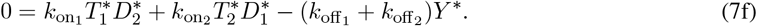

We will start with establishing several properties of nonnegative steady states of the system (1).

#### Lemma 1.

*Assume that the rate constants* 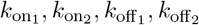 *are positive. Then, for every solution of the system* (7) *with nonnegative coordinates, the following hold:*

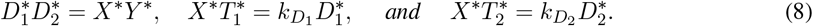

Proof. The projection of the solution set on the coordinates 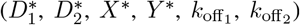 -coordinates (obtained by performing elimination with Gröbner bases [51, Chapter 2, §1]; the Maple code can be found in the supplementary materials [48]) yields the following equation:

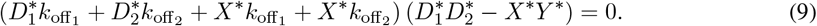

If the left bracket vanishes, then, by the positivity of the rate constants and nonnegativity of the solution, we deduce 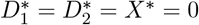, so 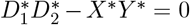. Otherwise, the right bracket must vanish, so the same equality holds. This gives us the first equality in (8). Adding this equation to (7) and computing projections (again with Gröbner bases) to 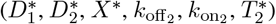 - and 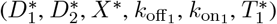 -coordinates, respectively, we obtain that:

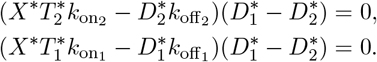

If 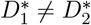, the above equalities imply the last two equations from (8).

Now we assume that 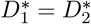, and add this equation to the system. By projecting on 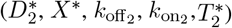 - and 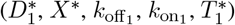-coordinates, respectively, we obtain

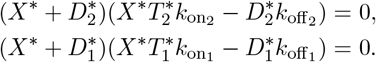

In both equations above, vanishing of the left bracket would imply the vanishing of the right, so the right one always vanishes. This yields the two remaining equations of (8) and concludes the proof. □

The three-body model (1) has three conservation laws:

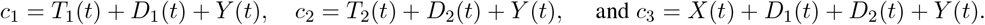

The set of nonegative states with the same values of *c*_1_, *c*_2_, *c*_3_ is referred to as *stoichiometric class*. Based on the initial conditions *D*_1_(0) = *D*_2_(0) = *Y* (0) = 0, we have *c*_1_ = *T*_1_(0), *c*_2_ = *T*_2_(0), and *c*_3_ = *X*(0). In the notation of Section 2.5, the lemma below establishes that, for the set of trajectories 𝒯 with these initial conditions and satisfying *c*_1_, *c*_2_, *c*_3_ *>* 0, the question of identifiability from steady states is well-posed.

#### Lemma 2

(cf. [52, p. 11]). *Assume that the rate constants* 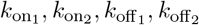 *are positive. Then, in each stoichiometric class (i*.*e*., *for every positive values of c*_1_, *c*_2_, *c*_3_*), there exists exactly one positive steady state which is globally asymptotically stable. Furthermore, its* 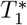 *and* 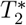 *-coordinates are the unique positive roots of*

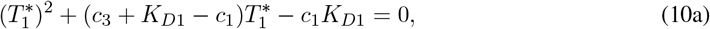

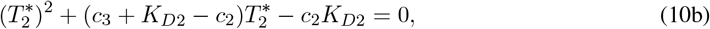

*and the X*^∗^*-coordinate satisfies*

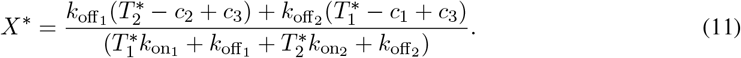

*Proof*. The existence and uniqueness of globally asymptotically stable equilibrium was established in [52, p. 11]. We recall the argument here. The system (4) can be represented by a chemical reaction network

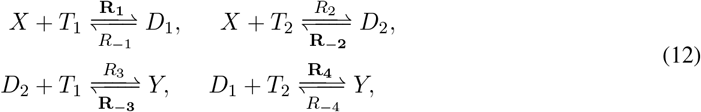

With the rates 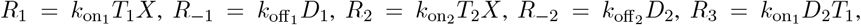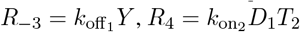, and 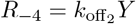.

The subnetwork of (12) consisting of the boldfaced reactions *R*_1_, *R*_−2_, *R*_−3_, *R*_4_ forms an M-network (see [52, p. 29]) and satisfies the requirements of [52, Theorem 4]: each species participates in exactly one reaction and appears as a product of exactly one reaction, this is easy to see from the Petri net representation of the subnetwork on Figure 8. Therefore, by [52, Theorem 4] the function max ℛ(*x*) − min ℛ(*x*), where

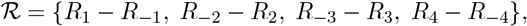

is a convex robust Lyapunov function. The existence of such function implies the existence, uniqueness, and global stability of an equilibrium in every stoichiometric class.

**Figure 8:**
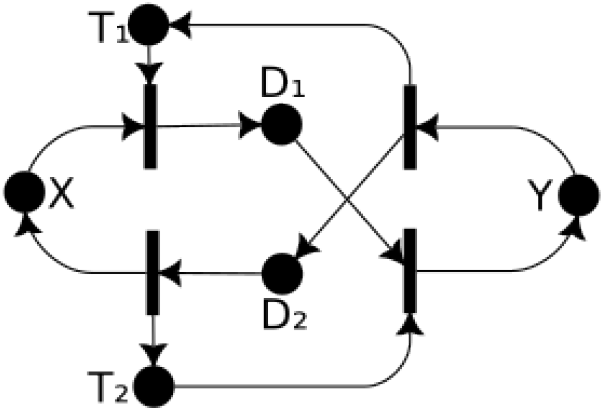
Petri-net representation of the chosen irreversible subnetwork of (12)

We will now establish formulas (10) and (11). We augment the system (7) with the equations:

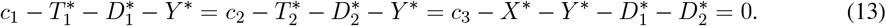

Given the obtained system of nine equations, we next compute the projections (again, using Gröbner bases; the Maple code can be found in the supplementary materials [48]) of the solution set onto the 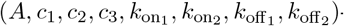 -coordinates, where *A* is taken to be 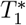 or 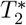. We find that these projections satisfy precisely (10). Consider (10a) as a quadratic equation in 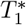. The product of the roots is equal to −*c*_2_*K*_*D*2_ *<* 0, from which we conclude that there exists exactly one positive root, so it must be the 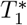 -coordinate of the unique steady state. The same applies to 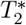.

Now we compute the projection onto the 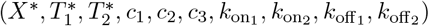-coordinates and find a relation:

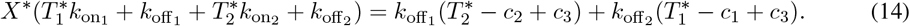

The positivity of 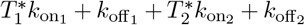 implies that we can express *X*^∗^ uniquely and obtain (11). □

In other words, Lemma 2 shows that 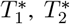, and *X*^∗^ can be expressed as algebraic functions of 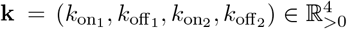 and *c*_1_, *c*_2_, *c*_3_ ∈ ℝ_>0_. We will denote these functions by the same letter and write, for example 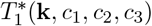 for the positive root of (10a).

Next, we will consider several (two or three) experiments with the same *c*_1_ = *T*_1_(0) and *c*_2_ = *T*_2_(0) but varying *c*_3_ = *X*(0) (cf. Example 5). The next proposition establishes that the vector 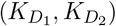 is identifiable up to at most two possible values from the steady state data of two such experiments.

#### Proposition 1.

*Consider two vectors* 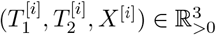 *for i* = 1, 2. *Assume that*

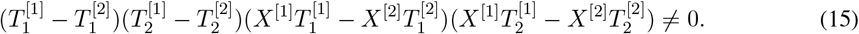

*Then, there exists a subset* 𝒦 ⊂ ℝ_>0_ *of cardinality at most two such that, for every* 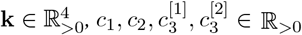 *satisfying*

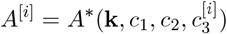

*for every* 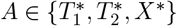 *and i* = 1, 2, *we have* 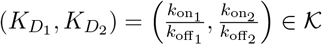.

*Furthermore, for every* 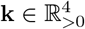 *and* 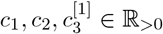, *there are only finitely many values* 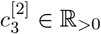 *such that* (15) *does not hold for* 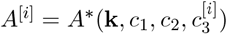, *where* 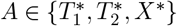 *and i* = 1, 2.

*Proof*. Consider two vectors 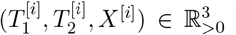 for *i* = 1, 2 and 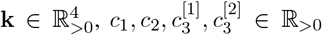 such that 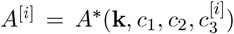 for every 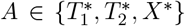 and *i* = 1, 2. We use (8) and equalities 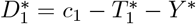 and 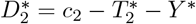 to write the following polynomial system:

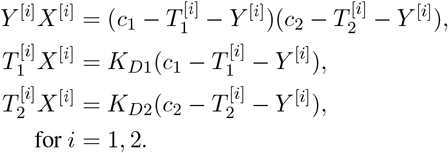

By computing the projection of the solution set of this system to the 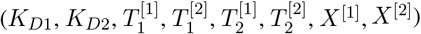-coordinates (using Gröbner bases), we find expressions of the form:

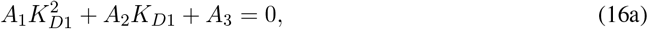

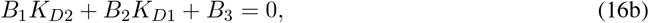

where *A*_1_, *A*_2_, *A*_3_, *B*_1_, *B*_2_, *B*_3_ are polynomials from 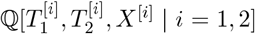. Furthermore, *A*_1_ and *B*_1_ factor as follows:

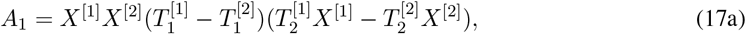

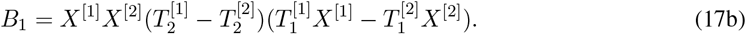

Therefore, if (15) holds, *A*_1_ and *B*_1_ do not vanish. Then 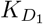 must be one of the roots of (16a) and then 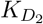 is uniquely determined from (16b). Since the coefficients of the equations (16a) and (16b) depend only on 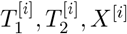 for *i* = 1, 2, these two solutions form the set 𝒦 from the statement of the proposition.

In order to prove the second statement of the proposition, we fix 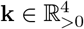 and 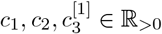 and consider 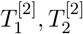 and *X*^[2]^ as functions in 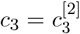. By Lemma 2, these functions are algebraic. We notice that (15) is equivalent to inequalities:

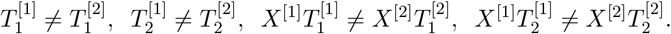

Since a nonconstant univariate algebraic function takes each value only finitely many times, it is sufficient to prove that the functions

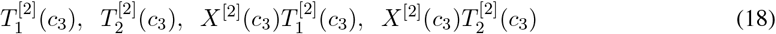

are nonconstant. To this end, we use (10a) and (10b) to write the Taylor series for 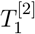 and 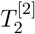 at *c*_3_ = 0:

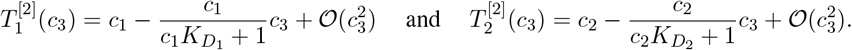

By plugging these to (11), we obtain an expansion for *X*^[2]^:

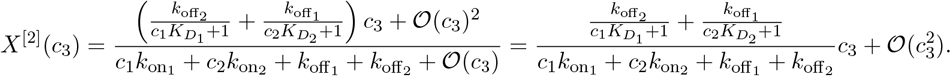

From these expansions, we see that indeed none of the functions (18) is constant. This finished the proof of the second claim of the proposition. □

The following theorem establishes that, using the terminology of Section 2.5, the vector 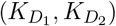 is globally identifiable from the steady state data for three generic experiments with the same values of *c*_1_, *c*_2_ but varying value of *c*_3_.

#### Theorem 1

(Unique identifiability from three experiments). *Consider three vectors* 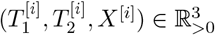 *for i* = 1, 2, 3. *Assume that*

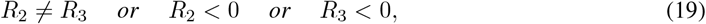

*where*

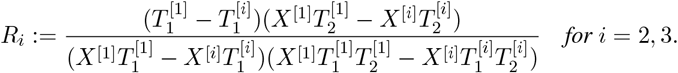

*Then there exists a vector* 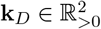 *such that for every* 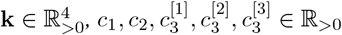 *satisfying*

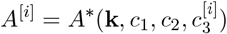

*for every* 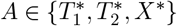 *and i* = 1, 2, 3, *we have* 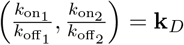.

*Furthermore, for every* 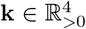 *and* 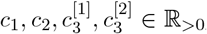, *there are only finitely many values* 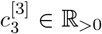 *such that* (19) *does not hold for* 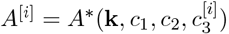, *where* 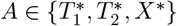 *and i* = 1, 2, 3.

*Proof*. The proof strategy will be similar to the one for Proposition 1. Consider two vectors 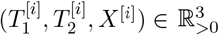 for *i* = 1, 2, 3 and 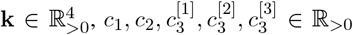 such that 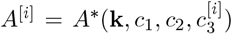 for every 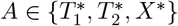 and *i* = 1, 2, 3.

Consider the relations (16a) and (16b) obtained in the proof of Proposition 1. Recall that (17a):

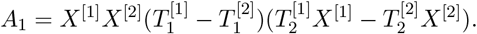

Furthermore, we have

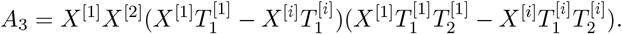

Therefore,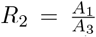. If *R*_2_ *<* 0, then (16a) has exactly one positive root. In this case, 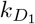 would be uniquely determined, and then 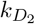 would be uniquely determined from (16b). The same argument applies to *R*_3_ *<* 0 due to the symmetry between the second and the third experiment.

In order to take into account the third experiment, we apply the derivation of (16a) and (16b) from the proof of Proposition 1 to the first and third vectors instead of the first and the second. This way, 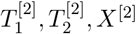in (16a) will be replaced with 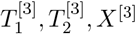, and the resulting relation will also be true. This will yield one more quadratic equation for 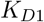, which we will denote by 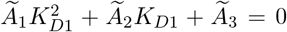. Since 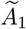 and 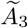 are obtained from *A*_1_ and *A*_3_ by replacing 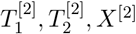 with 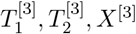, we have 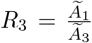. If 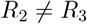, then the quadratic equations (16a) and 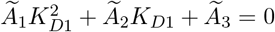 are not proportional, so they have at most one common root thus leaving at most one possible value for 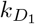.

In order to prove the second part of the theorem, we fix 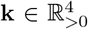 and 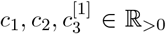, and consider the function

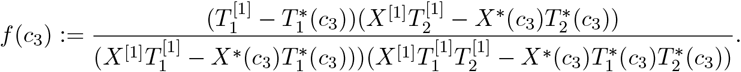

Then we have 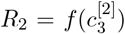 and 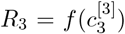. By Lemma 2, *f*(*c*_3_) is an algebraic function. Therefore, if it is a nonconstant algebraic function, it takes each value only finitely many times, so the equality *R*_2_ = *R*_3_ will be true only for finitely many values of 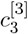 as desired. Now assume that *f*(*c*_3_) is a constant function. Then 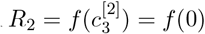. Using *X*^∗^(0) = 0 and 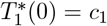, we find that

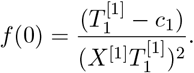

Since 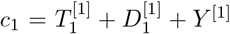, we have 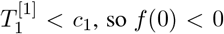. This implies *R*_2_ *<* 0, so (19) is fulfilled independently of the value of 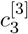. □

We will illustrate the above argument with a numerical example. We will first take the initial concentrations and parameter values the same as in the example in the introduction:

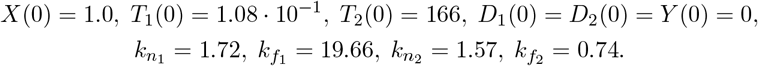

Through numerical simulations, we compute the values of *X, T*_1_, *T*_2_ at the steady state: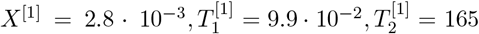. Next, we simulate one more experiment with all the initial conditions staying the same except for *X*(0) = 1.1. The new steady state is 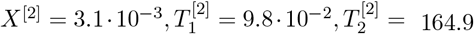. Plugging these values into equations (16), we find two solutions for 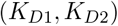 one of which is equal to the true values, and the other one is negative and can therefore be discarded. So, in this case, two experiments are sufficient to find the unique parameter values. Even if the second solution of (16) was positive, it could be discarded by performing one more experiment with different *X*(0), and choosing the solution of (16) appearing among the solutions of (16) for the first pair of experiments.

The details of all the symbolic and numeric calculations from this section can be found in the *identifiability* folder of the repository [48] with the supplementary code for the paper.

## 4 Discussion

Although each of the TCEs presented in this study was designed for a different cancer cell target, the initial concentration of the targets on cancer cells is assumed to be equally expressed on the cancer cell surface to visualize the quantitative differences between the bell-shapes in Figure 5.

From the three-body model perspective, a promising TCE antibody is the one that creates the largest trimer concentration with the minimal initial concentration of the antibody at the site of action. Additionally, to overcome natural intra-tumoral and inter-patient variability, it is favorable to have a TCE antibody with minimal variation for different ratios of the initial concentrations of the targets. So, a TCE antibody molecule is effective at the site of action if it results in: 1) a relatively higher peak of the trimer concentration at the site of action, 2) a relatively lower exposure in the circulating compartment to avoid toxicity issues, 3) lower affinity toward CD3 to prevent drug sing in the non-tumor compartments, and 4) robust characteristics of the bell-shape for different range of target concentrations, to have the highest predictability for different initial conditions across patients and tumors.

Among the molecules presented in Figure 6, it can be observed that the 7370, and PF-06671008 molecules are projected to have the largest concentrations of trimers in TME with minimal initial target concentrations. On the other hand, Solitomab creates a stable width of the peak for a wide range of ratios between the initial concentrations of the targets. It is apparent that stability across different initial concentrations of the targets increases the consistency of the data in clinical research.

In addition to the TCE molecules presented in Table 1, other molecules are discussed in detail in the literature. PF-06863135 [53] (150 kDa), and AMG420 [11] (54 kDa) are both designed to target B-Cell Maturation Antigen (BCMA) for patients diagnosed with multiple myeloma. Both of the molecules have similar dissociation constants to CD3, but different dissociation constants to BCMA, 0.1 and 0.04 nM respectively. The role of binding kinetics to BCMA is discussed as an important factor in the distribution of the molecule toward different tissues in [11]. Also, the authors of [54] discussed the design and bio-distribution of AMG211 (55 kDa), a CD3/CEA TCE molecule for patients with advanced gastrointestinal adenocarcinomas. AMG211 has a significantly high dissociation constant to CD3 310 nM in comparison with its dissociation constant toward CEA 5.5 nM. The biodistributions of these molecules are explained by a computational model in [55].

A molecule with a bell-shaped efficacy curve will have an important property: its efficacy will be maximized at the concentrations that correspond to the peak of trimer formation (Figure 1(b), yellow thick line), and will be low at the concentrations that are significantly less or significantly higher than that. This provides an opportunity to assess the predictions in the following way: with the understanding of PK properties of the desired compound, one can calculate doses (using standard PK-PD modeling methodology) that will achieve steady state concentrations in the tumor microenvironment below, at, or beyond the drug concentrations predicted to maximize efficacy. One can then design a pre-clinical experiment, where animals receive drug doses that will achieve these concentrations, and assess, whether efficacy is optimized at the intermediate “optimal” concentration. It should be noted that because it takes time for a drug to accumulate in the TME, it may take time to see the difference between the dosing regimens, with the higher-dose regimen achieving higher efficacy short term (it will reach the “optimal” drug concentration sooner) but will then lose efficacy in the longer term, since drug will continue accumulating, and therefore will soon move into the suboptimal range on the right of the efficacy curve.

Interestingly, if one were to evaluate the dosing regimens of some of the approved TCEs, one would observe that epcrotitamab (anti-CD3/anti-CD20) protocol dictates increasing spaces between doses, with the drug given weekly during the first three treatment cycles, then given at 48 mg every 2 weeks between 4th-9th treatment cycles, and finally every 4 weeks after the 10th treatment cycle. Glofitamab, another highly efficacious anti-CD3/anti-CD20 antibody, is given at 30 mg every 2 weeks for the first treatment cycle, and then every 4 weeks for subsequent cycles. Mosunetuzumab starts with weekly dosing, with 60 mg given every 3 weeks for the second treatment cycle, and then lowering the dose to 30mg for subsequent treatment cycles. All of these (increasing spacing between doses or lowering dosing for later treatment cycles) could be potential mitigation strategies to ensure that the drug concentration remains in the “optimal” zone.

The simplified three-body model provides valuable initial insights, but it may not fully capture the complexity of TCE interactions *in vivo*. Tailored version of this model for each target to incorporate factors such as T cell dynamics, tumor heterogeneity, and drug disposition would improve the model’s predictive accuracy for clinical applications for future programs.

A common safety concern for TCEs is cytokine release syndrome (CRS), which arises from excessive immune activation and leads to a surge in inflammatory cytokines with potentially severe systemic effects. While the present study focuses on characterizing dose-response relationships with respect to efficacy—using maximum trimer concentration as a key feature—understanding the full therapeutic window will ultimately require integrating both efficacy and safety considerations.

In particular, the bell-shaped nature of the dose-response curve suggests that exceeding a certain concentration could result not only in reduced efficacy but also in heightened toxicity risk. Thus, future modeling efforts should aim to incorporate safety endpoints, such as cytokine profiles or clinical markers of CRS, to better inform dose selection. A recent review of clinical dose optimization strategies for T-cell-enhanced therapies highlights various dosing approaches and emphasizes the importance of customized dosing regimens via mathematical modeling in the future [56].

Additionally, while our study emphasizes the peak of the bell-shaped curve as a marker of maximum efficacy, we recognize that this alone is not sufficient to fully characterize the dose-response relationship. We complement our peak-focused sensitivity analysis with numerical comparisons of the curve width (e.g., Figure 6), which offer insight into the robustness and breadth of the therapeutic window. A more comprehensive modeling framework—including toxicity metrics and safety thresholds—will be necessary to support clinical dose optimization, and this remains an important direction for future work.

The proposed analysis can be used to further assess the optimal properties for the design of CD3-based bispecific T cell engagers for a variety of scenarios and targets, to hopefully expand the applicability of this modality to a larger number of indications.

## 5 Acknowledgments

The authors thank Anup Zutshi, Kumpal Madrasi, and Abed Alnaif for the discussions and constructive feedback. The biodistribution model is significantly enhanced by the comments received from Thang Ho.

## 6 Author Contributions

MS, IK, GP, and EDS wrote the manuscript; MS, IK, and EDS designed the research; MS and GP performed the research; MS, IK, and EDS analyzed the data.

## Footnotes

1 The three-body model (1) in the required format can be found in the github repository at https://github.com/mahdiarsadeghi/tce/blob/main/identifiability/00_time_course.md

## Notes

**Conflict of interest:** The authors declare that the research was conducted in the absence of any commercial or financial relationships that could be construed as a potential conflict of interest.

**Funding:** MS and EDS were partially supported by NSF grant DMS-2052455, and AFOSR grant FA-9550-21-1-0289. GP was partially supported by NSF grants DMS-1853482, DMS-1760448, and DMS-1853650 and by the French ANR-22-CE48-0008 OCCAM project. MS was a summer intern, and IK is an employee, of EMD Serono Inc., a US subsidiary of Merck KGaA. MS is not currently affiliated with EMD Serono Inc. Views expressed in this paper do not necessarily represent the views of EMD Serono Inc.

https://github.com/mahdiarsadeghi/bites

### Competing Interest Statement

The authors have declared no competing interest.

